# Signatures of positive selection and local adaptation to urbanization in white-footed mice (*Peromyscus leucopus*)

**DOI:** 10.1101/038141

**Authors:** Stephen E. Harris, Jason Munshi-South

## Abstract

Urbanization significantly alters natural ecosystems and has accelerated globally. Urban wildlife populations are often highly fragmented by human infrastructure, and isolated populations may adapt in response to local urban pressures. However, relatively few studies have identified genomic signatures of adaptation in urban animals. We used a landscape genomics approach to examine signatures of selection in urban populations of white-footed mice (*Peromyscus leucopus*) in New York City. We analyzed 154,770 SNPs identified from transcriptome data from 48 *P. leucopus* individuals from three urban and three rural populations, and used outlier tests to identify evidence of urban adaptation. We accounted for demography by simulating a neutral SNP dataset under an inferred demographic history as a null model for outlier analysis. We also tested whether candidate genes were associated with environmental variables related to urbanization. In total, we detected 381 outlier loci and after stringent filtering, identified and annotated 19 candidate loci. Many of the candidate genes were involved in metabolic processes, and have well-established roles in metabolizing lipids and carbohydrates. Our results indicate that white-footed mice in NYC are adapting at the biomolecular level to local selective pressures in urban habitats. Annotation of outlier loci suggest selection is acting on metabolic pathways in urban populations, likely related to novel diets in cities that differ from diets in less disturbed areas.

## INTRODUCTION

Urban habitats are one of the fastest growing and most rapidly changing environments around the world. While urbanization has been traditionally viewed as a driver of declining habitat quality in and around cities, there is growing interest in the idea that urban areas represent novel environments with unique selective pressures (Donihue & Lambert 2015). The recently developed but burgeoning field of urban evolutionary biology aims to determine how urbanization leads to evolutionary change through mutation, genetic drift, gene flow, and natural selection in urban populations.

The ecological changes that occur within cities are likely to have many evolutionary implications. Human infrastructure causes habitat loss and fragmentation and changes resource availability, novel species interactions occur because human movements and commerce introduce a diverse array of nonnative species, and human activity increases exposure to chemical, light, and noise pollution (McKinney 2002; Chace & Walsh 2004; Shochat *et al.* 2006; Sih *et al.* 2011). These changes lead to unique pressures in novel urban habitats that may rapidly drive evolutionary change over short timescales. Increased genetic drift in relatively isolated urban populations, genetic differentiation between populations with restricted gene flow from urban infrastructure, or allele frequency shifts due to local urban adaptation, are all likely outcomes of evolution in cities (Munshi-South 2012; Merilä & Hendry 2014; Donihue & Lambert 2015).

Urban populations are potentially excellent systems for examining how species respond to anthropogenic environmental change, what genes and traits are involved, and how quickly populations locally adapt to changing environments. Local adaptation is a common phenomenon in nature (Stinchcombe & Hoekstra 2008; Bonin 2008; Linnen *et al.* 2009; Hohenlohe *et al.*2010a; Turner *et al.* 2010; Ellison *et al.* 2011; De Wit & Palumbi 2013), and often results from the operation of selection on standing genetic variation as opposed to novel mutations over relatively short time scales (Barrett & Schluter 2008; Stapley *et al.* 2010). Additionally, the quantitative traits involved in local adaptation may involve many genes of small effect working to produce the desired phenotype (Orr 2005; Rockman 2012), and these ecologically relevant but non-conspicuous phenotypes are predicted to be those most involved in urban adaptation (Sih *et al.* 2011). However, traits with relatively simple genetic architecture may also be under selection in urban environments (Thompson *et al.* 2016). Investigating the genetic basis of local adaptation has provided insight into a variety of evolutionary processes including speciation, maintenance of genetic diversity, range expansion, and species responses to changing environments (Savolainen *et al.* 2013; Tiffin & Ross-Ibarra 2014), and holds great promise for understanding adaptive evolution in response to urbanization.

Landscape genomics has recently produced a number of approaches for studying local adaptation. This field is defined by the spatially explicit study of genomic variation (Sork *et al.* 2013) that seeks to identify environmental variables influencing adaptive genomic variation (Rellstab *et al.* 2015). Landscape genomics, and more specifically genotype-by-environment analyses (GEA), can successfully identify associations between urban environmental variables and allele frequencies that indicate adaptation to local urban conditions. These approaches can also help to untangle the interactions between neutral demographic processes and selection (Rellstab *et al.* 2017). Urban populations are influenced by both genetic drift through founder effects and barriers to gene flow, and selection acting on genetic variation linked to increased fitness in urban settings.

A small but growing number of studies have documented how populations may locally adapt to urban selective pressures through changes in allele frequencies and / or undergo directional shifts in phenotypic traits. Yeh (2004) reported that sexually-selected tail coloration in Juncos (*Junco hyemalis*) was rapidly evolving in urban populations compared to rural ones. European Blackbirds (*Turdus merula*) exhibit evidence of selection on genes underlying anxiety behavior in newly established populations across multiple cities (Partecke *et al.* 2006; Mueller *et al.* 2013). Cheptou *et al.* (2008) reported that a weed (*Crepis sancta)* in urban vegetation plots surrounded by paved surfaces showed heritable changes in seed morphology and dispersal. Reduced snow cover in urban areas leads to colder minimum ground temperatures and Thompson *et al.* (2016) found parallel adaptive evolution in urban white clover (*Trifolium repens*) populations that had increased freezing tolerance. Several studies have also found likely adaptive genetic and morphological changes in urban mammal populations. Suggestive of urban adaptation, a specific mitochondrial genotype rose to fixation in white-footed mice (*Peromyscus leucopus)* populations in Chicago along with morphological changes to skull shape after urbanization (Pergams & Lacy 2008). In urban areas of Italy, Kuhl’s pipistrelle (*Pipistrellus kuhlii*) bat populations had significantly larger bodies and longer skulls than natural populations, suggesting urban adaption to a novel diet introduced when artificial illumination attracted an increased number of large hard-bodied moths (Tomassini *et al.* 2014).

Few studies in urban evolutionary biology have been able to measure phenotypic changes, definitively link them to genetic changes, and establish fitness benefits to demonstrate evolutionary adaptation. One exception are urban killifish (*Fundulus heteroclitus*), where selective pressure from polychlorinated biphenyls (PCBs) has led to the evolution of PCB tolerance in urban populations (Whitehead *et al.* 2010; Reid *et al.* 2016). Adaptation to PCB pollution was also reported in tomcod (*Microgadus tomcod)* in the Hudson River through a deletion that similarly increases tolerance to PCBs (Wirgin *et al.* 2011). Urban adaptation has also been confirmed in the well-known peppered moth (*Biston betularia*) system. Recent evidence suggests that the industrial melanism trait in this species is linked to an insertion of a transposable element in the *cortex* gene in the early 1800s that spread throughout the population in response to industrial airborne pollution (Hof *et al.* 2016). The study of additional systems will likely identify a complex array of adaptive evolutionary responses in cities (Whitehead *et al.* 2017).

Here we examined signatures of selection in isolated urban populations of white-footed mice, *Peromyscus leucopus*, in New York City (NYC) using a landscape genomics approach. *Peromyscus* spp. (Rodentia, Cricetidae) are a group of abundant small mammals found across much of North and Central America. They live in a diverse array of habitats that exposes them to a variety of selective pressures, and thus multiple *Peromyscus* spp. have become model systems for studies examining ecology, evolution, and physiology in natural populations (Munshi-South & Richardson 2017). There is also evidence that *Peromyscus* spp. readily adapt to environmental change (Storz *et al.* 2007, 2009, 2010; Mullen & Hoekstra 2008; Linnen *et al.* 2009; Weber *et al.* 2013; Natarajan *et al.* 2013; Munshi-South & Richardson 2017), making them good subjects for the study of local adaptation. White-footed mice are one of the few native mammals that thrive in extremely small, fragmented urban forests in North America (Pergams & Lacy 2008; Rogic *et al.* 2013; Munshi-South & Nagy 2014), and tend to be found at higher densities in urban vs. rural patches due to a thick understory providing abundant food resources and exclusion of major predators and competitors (Rytwinski & Fahrig 2007). Increased density may also be due to limited *P. leucopus* dispersal between urban sites. Munshi-South (2012) Hfound barriers to dispersal between isolated NYC parks, with migrants only moving through significantly vegetated corridors throughout the city. There is also substantial genetic structure between NYC parks as measured by microsatellites (Munshi-South & Kharchenko 2010), genome-wide SNPs (Munshi-South *et al.* 2016) and demographic modeling (Harris *et al.* 2016). We have also previously identified signatures of selection in urban populations of NYC white-footed mice (Harris *et al.* 2013), though we used smaller datasets and more limited approaches than presented here.

In the current study, we examined SNPs generated from individual transcriptome sequencing for *P. leucopus* from three urban sites in NYC and three rural sites from the surrounding area. We generated a large SNP dataset and produced estimates of nucleotide diversity (π, Tajima 1983), Tajima’s *D* (Tajima 1989), and *F*_*ST*_ (Wright 1951) to generate persite estimates and identify loci that deviate from neutral expectations. We then used a variety of genome scan methods and outlier tests to identify genes subject to selection in an urban setting. Our approach identified population differentiation, shifts in allele frequencies, and associations between alleles and environmental variables. However, neutral demographic processes such as population bottlenecks can produce signatures of genetic variation similar to those produced by selection (Oleksyk *et al.* 2010; Li *et al.* 2012). We accounted for this possibility by incorporating a simulated neutral SNP dataset from an inferred demographic history (Harris *et al.* 2016) directly into our null model for identifying outliers (Excoffier *et al.* 2009; Gutenkunst *et al.* 2009; Li *et al.* 2012; Vitti *et al.* 2013; Lotterhos & Whitlock 2015).

The three specific aims of this study were the following: 1. identify candidate genes exhibiting signatures of selection in NYC populations of white-footed mice using a variety of genome scan methods and outlier tests; 2. distinguish genetic outliers resulting from selection rather than demography by incorporating demographic histories of white-footed mice in NYC into null models of genome scans; and 3. identify genes that are statistically associated with environmental variables representative of urbanization using landscape genomic approaches.

## MATERIALS AND METHODS

### Sampling, library preparation, and transcriptome assembly

We trapped and collected white-footed mice from 2010 – 2012. For full details on sampling, transcriptome sequencing, assembly and SNP calling, see Harris *et al.* 2013, 2015. In brief, we randomly chose eight adult white-footed mice (equal numbers of males and females) from each of six sampling locations (N = 48 total) representative of urban and rural habitats and with minimal within-site genetic structure (Fig. 1) (Harris *et al.* 2013, 2015). Three sampling sites were within NYC parks: Central Park in Manhattan (CP), New York Botanical Gardens in the Bronx (NYBG), and Flushing Meadows—Willow Lake in Queens (FM). These sites represented urban habitats surrounded by high levels of impervious surface cover and high human population density, as previously quantified in Munshi-South et al. (2016). The remaining three sites occurred ∽100 km outside of NYC in rural, undisturbed habitat representative of natural environments for *P. leucopus*. High Point State Park is in the Kittatinny Mountains in New Jersey (HIP), Clarence Fahnestock State Park is located in the Hudson Highlands in New York (CFP), and Brookhaven and Wildwood State Parks occur on the northeastern end of Long Island, New York (BHWWP). Total RNA was extracted separately from livers stored in RNA later for each of the 48 mice, treated with DNase, enriched through ribosomal RNA depletion, fragmented, reverse transcribed, amplified and tagged with a unique barcode, and sequenced in four lanes of one SOLiD 5500XL run (Harris *et al.* 2015). We called SNPs with the Genome Analysis Toolkit (GATK version 2.8) pipeline using a Bayesian genotype likelihood model (DePristo *et al.* 2011). In order to call a SNP, we required it to occur in at least five individuals, have a nucleotide quality (q-score) ≥ 30, exhibit no strand bias (FS ≥ 35), and to come from only uniquely mapped reads. We also required SNPs to have an overall depth ≥ 10X and ≤ 350X (to account for paralogous sequences), a minor allele frequency (MAF) ≥ 0.025, and removed SNPs where every individual was heterozygous.

**Figure 1.**
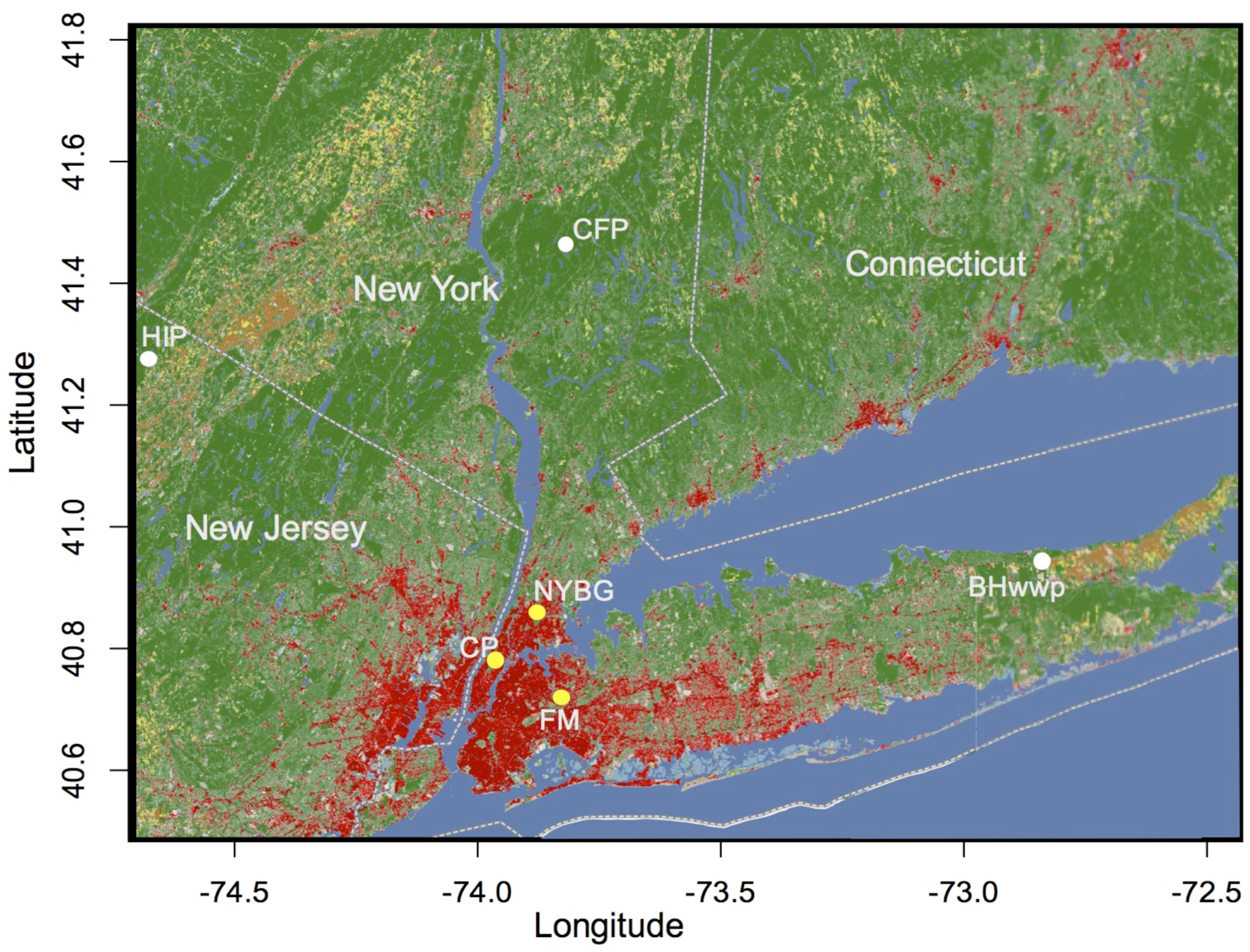
Map of sample localities in the NYC metropolitan area. Sites in yellow are urban parks within New York City, CP = Central Park; FM = Flushing Meadows—Willow Lake; NYBG = New York Botanical Gardens. Sites in white are rural parks, BHwwp = Brookhaven and Wildwood State Park; CFP = Clarence Fahnestock State Park; HIP = High Point State Park. The map includes data from the National Land Cover Database. All non-green colors are shaded according to land use. Yellows and browns equal cultivated land and reds represent developed areas (Darker red = increased development). Green colors are shaded according to canopy cover (Darker green = increased canopy cover) and come from the 2011 National Land Cover Canopy database. Full legends for the colors are shown in Figure S1.

### Summary statistics

SNP information was stored in a VCF (variant call format) file and summary statistics were calculated using vcftools 0.1.12b (Danecek *et al.* 2011). We calculated per-site nucleotide diversity (π), Tajima’s *D*, and *F*_*ST*_. We also calculated the statistics for each contig (per-site statistic summed across all SNPs per contig divided by total sites) and calculated the average estimate for each population, including all pairwise population comparisons for *F*_*ST.*_

### Scans for positive selection based on population differentiation

We used the *F*_*ST*_ based analysis implemented in BayeScan v. 2.1 (Foll & Gaggiotti 2008) to compare all six population-specific allele frequencies with global averages and identify outlier SNPs. BayeScan identifies loci that exhibit divergence between groups that is stronger than would be expected under neutral genetic processes. Based on a set of neutral allele frequencies under a Dirichlet distribution, BayeScan uses a Bayesian model to estimate the probability that a given locus has been subject to selection. To generate more realistic allele frequency distributions, we used BayeScan for independent coalescent simulations of SNP datasets based on a neutral demographic history inferred by Harris *et al.* (2016) specifically for each *P. leucopus* population. Using the coalescent-based fastsimcoal2 software (Excoffier *et al.* 2013), we generated 100 sets of 100,000 SNPs each for every population in this study from a three population isolation-with-migration model using parameter estimates for divergence time, effective population size, migration rate, and population size change previously inferred in Harris *et al.* (2016). In short, the model represented a deep split between an ancestral population into Long Island, NY and the mainland (including Manhattan) 29,440 generations before present (GBP). A third population (representing the sampling sites in this study) later became isolated 746 GBP. Urban populations were also modeled to include a population size change event at the time of divergence. BayeScan was run independently on each of the 100 simulated datasets from fastsimcoal2 using default parameters to generate a null distribution of BayeScan statistics.

BayeScan was then run on the observed SNP dataset using default parameters. We performed several different analyses including a global analysis, one with two populations representing urban and rural groups, and finally analyses on all sampling site pairwise comparisons. We retained outlier SNPs with a q-value ≤ 0.1 (leading to a FDR of ≤ 0.1) and with a posterior odds probability from BayeScan higher than for any value calculated from the simulated dataset. BayeScan also calculates alpha (α), a locus specific F_st_ coefficient, where a positive value suggests diversifying selection and a negative value suggests balancing or purifying selection. There were no SNPs with negative α values.

For comparison to BayeScan results, we used a related method, BayPass (Gautier 2015), that identifies loci subject to selection based on allele frequency patterns that deviate from neutral expectations. We ran BayPass using default parameters under the auxillary covariate (AUX) model, and simulated pseudo-observed datasets (PODs) under the Inference Model in Baypass as suggested by Gautier (2015) to calibrate neutral distributions for XtX. BayPass uses the XtX statistic to identify adaptive divergence. SNPs with XtX estimates greater than the 95% threshold determined from PODs were identified as resulting from adaptive divergence.

### Analysis for selective sweeps

We also identified outlier regions when the observed SFS showed an excess of low frequency and high frequency minor alleles, a signal indicative of a recent selective sweep. The composite likelihood ratio (CLR) statistic is used to identify regions where the observed SFS matches the expected SFS generated from a selective sweep (Kim & Stephan 2002; Nielsen *et al.* 2005; Pavlidis *et al.* 2010). We calculated the CLR along sliding windows across the transcriptome using the software program SweeD (Pavlidis *et al.* 2013). SweeD is an extension of Sweepfinder (Nielsen *et al.* 2005) that is optimized for large next generation sequencing (NGS) datasets. We lacked a genome to provide high-quality linkage information so SweeD was run separately for each population and on individual contigs. We used default parameters except for using a sliding window size of 200 bp and use of a folded SFS, as we lacked an outgroup to infer ancestral alleles. The window within each contig with the highest CLR score is considered the likely location of a selective sweep. Similar to the method used for BayeScan, statistical significance was established from a null distribution generated by running SweeD on SNP datasets simulated under the inferred demographic history for *P. leucopus* populations (Harris *et al.* 2016). SweeD does not inherently identify outlier regions. The CLR is computed using a selective sweep model on the observed data and then compared to a neutral model calibrated with a simulated background SFS. As before, we used 100 datasets with 100,000 SNPs each, simulated under the inferred neutral demographic history for white-footed mice in NYC. The CLR was calculated using SweeD for all simulated datasets. We identified outlier contigs if their CLR value was greater than any produced in neutral simulations. We also required outliers to fall within the top 0.01% of the CLR distribution for the observed SNPs.

### Genotype-environment association tests for environmental selection

We used the GEA approach of LFMM: Latent Factor Mixed Models (Frichot *et al.* 2013) to associate our full SNP dataset with potential environmental selection pressures. LFMM examines associations between environmental and genetic variation while accounting for the neutral genetic background and structure between populations (Frichot *et al.* 2013). We tested three environmental variables associated with urbanization: 1) percent impervious surface (i.e. surfaces such as roads, rooftops, and other human infrastructure that do not absorb water calculated from USGS National Land Cover Data) within a 2 km (the approximate lifetime dispersal distance of white-footed mice) buffer around each sampling site’s GPS coordinate, 2) human density within a two-kilometer buffer around each sampling site’s GPS coordinate (calculated from US Census blocks), and 3) categorization of each site as urban, within NYC limits, or rural, undeveloped state park outside city limits (Coded as 0 or 1 in LFMM).

Calculations were made in ArcGIS v10.1 (ESRI, Redlands, CA, USA) and were previously reported in Munshi-South *et al*. (2016). This previous analysis found that variables 1-2 were significantly associated with genome-wide variation in *P. leucopus* populations in the NYC metropolitan area. LFMM requires the user to define the number of latent factors, K, that describe population structure in the dataset. To identify the appropriate number of K latent factors, we performed a genetic PCA followed by a Tracy-Widom test to find the number of eigenvalues with *P* values ≤ 0.01 (Patterson *et al.* 2006; Frichot & François 2015). Based on this approach, we ran LFMM with default parameters except for K = 6, number of MCMC cycles = 100,000, and burn-in = 50,000. Using author recommendations, we calculated the median |z|- score from 10 replicate runs and then readjusted the p values. LFMM uses |z|- scores to report the probability of a SNP’s association with an environmental variable. Again, we controlled for FDR by using a q-value threshold of ≤ 0.1.

BayPass also includes an environmental analysis, so for comparison to LFMM we used the GEA test implemented in the BayPass AUX model that identifies genetic markers associated with population-specific covariates (Gautier 2015). For population covariates, we used the same environmental variables used in LFMM: site classification (i.e. urban or rural) as a binary covariate, human density, and impervious surface. We used the AUX model and again simulated pseudo-observed datasets (PODs) under the Inference Model to calibrate neutral distributions for Bayes Factors (BFs). BayPass uses BFs to associate SNPs with population specific covariates. SNPs with BF estimates greater than the 95% threshold determined from PODs were considered to be associated with population covariates. We further filtered associations by setting a cutoff for BF ≥ 20.

### Functional annotation of candidate genes

We used the gene annotation pipeline in Blast2GO (Conesa *et al.* 2005; Götz *et al.* 2008) to identify sequences from the NCBI non-redundant protein database that were homologous to our outlier contigs identified above. We then retrieved associated gene ontology (GO) terms. Blast2GO retrieves GO terms associated with BLASTX hits and uses the KEGG database to describe biochemical pathways linking different enzymes (Ogata *et al.* 1999; Kanehisa *et al.* 2014). For downstream enrichment analyses, we also used the Ensembl gene annotation system (Aken *et al.* 2016) to find homologous *Mus musculus* genes for each *P. leucopus* contig. We further interpreted the outlier gene lists using g:Profiler (Reimand *et al.* 2016) to identify gene ontology terms enriched in our outlier gene list compared to the fully annotated *Mus musculus* genome. We used the g:Profiler webserver and identified enriched terms associated with outlier genes using default parameters and the Benjamini–Hochberg correction for multiple comparisons with an adjusted p-value < 0.05. Finally, we used REViGO to cluster GO terms and summarize them in a subset of terms based on semantic similarity measures (Supek *et al.* 2011).

## RESULTS

### Genetic diversity statistics

**In total,** we identified 154,770 SNPs for investigating patterns of genetic variation and performing tests of selection. Urban populations had a 50% decrease in nucleotide diversity compared to the rural populations, but mean Tajima’s D values for rural parks were consistently higher than for urban parks (Table 1). The average nucleotide diversity for all three rural populations was 0.224 ± 0.034 SE, while the average for urban populations was only 0.112 ± 0.019 SE. The average Tajima’s *D* within populations did not show substantial differences between populations (Table 1). For all populations, Tajima’s *D* was slightly positive. Average pairwise *F*_*ST*_ were the lowest between rural populations (0.018 ± 0.364 SE, CFP – HIP Table S1) and highest between urban populations (0.110 ± 0.520 SE, CP – FM Table S1). These *F*_*ST*_ values were similar to *F*_*ST*_ estimated using genome-wide SNP datasets (Munshi-South *et al.* 2016).

**Table 1.** Summary population genomic statistics (mean ± standard error) for three urban and three rural populations of white-footed mice (*Peromyscus leucopus*) examined in this study.

| Population | Nucleotide diversity ($\pi$) | Tajima's $D$ | 945 |
| --- | --- | --- | --- |
| *Urban* |  |  | 946 |
| CP | 0.131 ±0.001 | 0.318 ±0.005 | |
| FM | 0.112 ±0.001 | 0.301 ±0.006 | 947 |
| NYBG | 0.092 ±0.001 | 0.280 ±0.006 | |
| *Rural* |  |  | 948 |
| BHwwp | 0.198 ±0.001 | 0.350 ±0.004 | 949 |
| CFP | 0.211 ±0.001 | 0.336 ±0.004 | |
| HIP | 0.263 ±0.001 | 0.349 ±0.004 | 950 |

### Outlier detection and environmental associations

We used BayeScan to identify 39 outlier SNPs exhibiting patterns of divergent selection between urban and rural populations (Fig. 2A, Table S2). There were no SNPs that exhibited signatures of balancing selection. *F*_*ST*_ values for outlier SNPs ranged from 0.21 – 0.33. BayeScan identified zero outlier SNPs in the simulated neutral dataset, and accordingly the 39 outlier SNPs from the observed data had q-values that were smaller than the most extreme values for the simulated data (q-value ≤ 0.6). We ran a similar test looking for patterns of divergence using BayPass. This analysis identified 56 SNPs that showed evidence of divergent selection (Table S2). We used PODs to estimate a null distribution and outlier SNPs had XtX values ≥ 8.35 (top 5% of the null distribution). There were 11 SNPs associated with diversifying selection in both the BayeScan and BayPass analyses.

**Figure 2.**
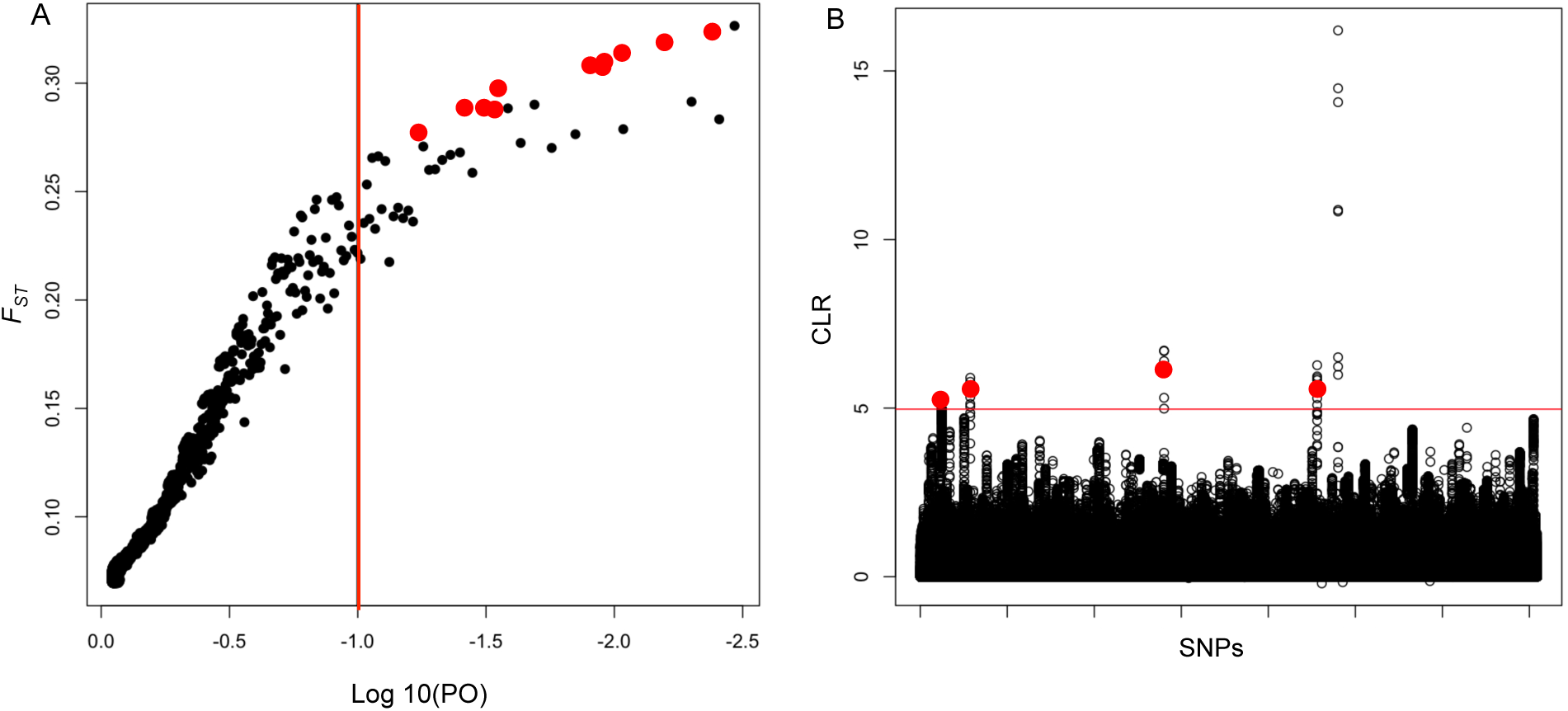
(a) BayeScan 2.1 plot of 154,770 SNPs genome scan analysis between urban and rural populations, including 48 individual white-footed mice from six NYC sampling sites. *F*_ST_ is on the vertical axis plotted against the log_10_ of the posterior odds (PO). The vertical red line indicates the cutoff (q-value = 0.1) used for identifying outlier SNPs. The markers on the right side of the vertical line show all outlier SNP candidates and the red circles represent the final accepted outlier SNPs from Table 2. (b) SweeD results with each of the 154,770 SNPs plotted from all 48 individuals. The Composite Likelihood Ratio (CLR) is plotted along the vertical access and each unfilled point represents an individual SNP. The x-axis has SNPs ordered by contig, but not by genomic position. The horizontal red line indicates the cutoff used for identifying outlier SNPs at *P* ≤ 0.0001. The red circles represent the final accepted outlier SNPs from Table 2.

**Table 2.**
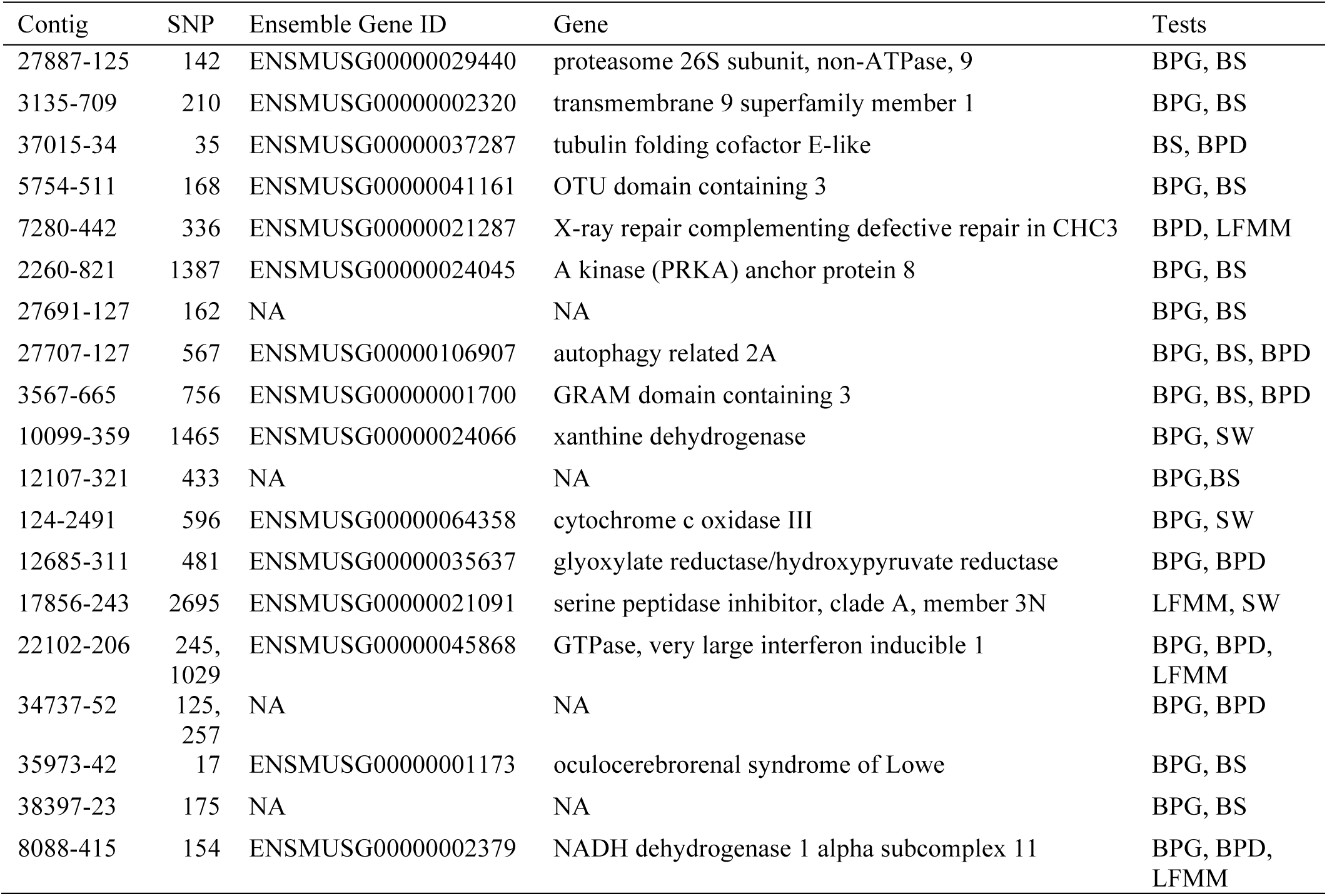
Outlier loci (*N* = 19) identified in at least one test for selection (BayeScan, BayPass, or SweeD) and one GEA test (LFMM or BayPass_GEA). SNP shows the position in contig containing the outlier loci. Tests show which tests identified the SNP as an outlier: BPG = BayPass_GEA; BPD = BayPass_Diversifying; BS = BayeScan; SW = SweeD; LFMM = LFMM.

To identify signatures of selective sweeps, we used the CLR statistic implemented in SweeD. We found that CLR scores in the top 5% of the simulated distribution were generally 2-3X lower than values in the top 5% of the observed dataset. We ran SweeD on observed SNPs within individual contigs and identified outliers by filtering for a CLR score ≥ 3.53 (the maximum CLR from simulated data). We also chose regions that fell within the top 0.01% of the observed distribution (Fig. 2B); all outliers had CLR scores ≥ 4.97. SweeD identified regions with SFS patterns that fit a selective sweep model in 45 contigs within urban populations (Table S2). There was no overlap between outlier SNPs identified by SweeD and BayeScan / BayPass.

There were 131 SNPs associated with at least one of three environmental variables tested using LFMM (Fig. 3A, Table S2). There was zero overlap with outliers identified from BayeScan and only one SNP that overlapped between SweeD and LFMM. Three SNPs identified in BayPass as outliers showing signatures of diversifying selection were also associated with environmental covariates in LFMM (Table S2). All three SNPs were within genes associated with human density around sampling sites and one was associated with all three environmental covariates. In an analysis similar to LFMM, we used BayPass to also associate environmental variables, called population covariates, with allele frequencies. There were 143 SNPs associated with at least one of the three environmental covariates tested using BayPass (Table S2). From these 143, five overlapped with those showing signatures of divergent selection in BayPass and eleven overlapped with outliers in BayeScan.

**Figure 3.**
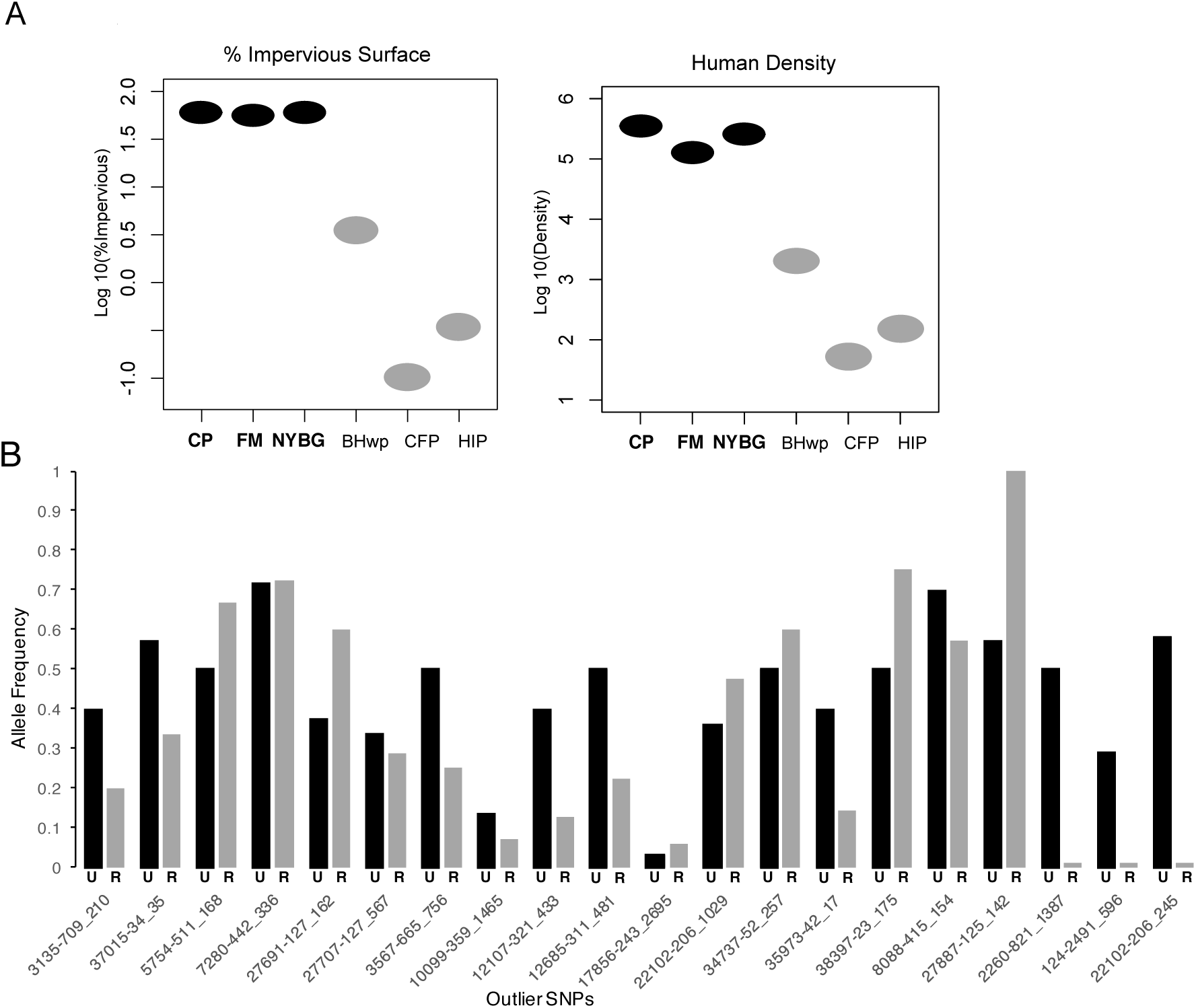
(a) Plot of urbanization metrics for all 6 sampling sites from NYC used in this study. Urban sampling sites are highlighted in bold on the horizontal axis and colored black. Rural sites are colored gray. The log10 value of % Impervious Surface and Human Density are plotted along the vertical axis and the oval represents the value for each sampling site. (b) Allele frequencies for candidate loci identified from both genome scans and GEA tests grouped by urban (U, black) or rural (R, gray) classification. The frequency of the outlier SNP within each type of population is plotted on the vertical axis. Each candidate loci is labeled with the contig and outlier SNP on the horizontal axis; see Table 2 for associated gene names.

Across all tests, SNPs identified as outliers or associated with environmental variables were found in 381 contigs. We filtered this list down to a subset of 19 contigs (Table 2) that are the most likely candidates for directional selection due to urban selective pressures. We required these filtered candidate contigs to show a signature of diversifying selection between urban and rural populations (BayScan or BayPass) or a signature of a selective sweep (SweeD), and they had to be associated with an environmental variable (human density around parks, impervious surface) as identified in GEA tests (LFMM or BayPass).

### Functional annotation

The full contig sequences containing outlier SNPs were obtained from the *P. leucopus* transcriptome (Harris *et al.* 2015) and used for functional annotation and analysis. We first tested the full set of 381 contigs identified by all outlier tests for overrepresented GO terms using g:Profiler. There were 260 overrepresented GO terms from the full outlier list (Table S3). We summarized this list using REViGO into 23 representative terms. The top representative term was lipid metabolism, followed by organic substance catabolism (Table S4). The list also includes lipid homeostasis and immune system processes.

We also looked for overrepresentation in the gene annotations associated with the filtered subset of 19 outliers and found related results (Table 3). There were 15 contigs homologous to known genes with functional annotation. Metabolic pathways were the most overrepresented group of gene ontology terms, and there were two biological functions associated with the most overrepresented GO terms from the full list. These included non-alcoholic fatty liver disease and regulation of protein kinase b (AKT) signaling.

**Table 3.**
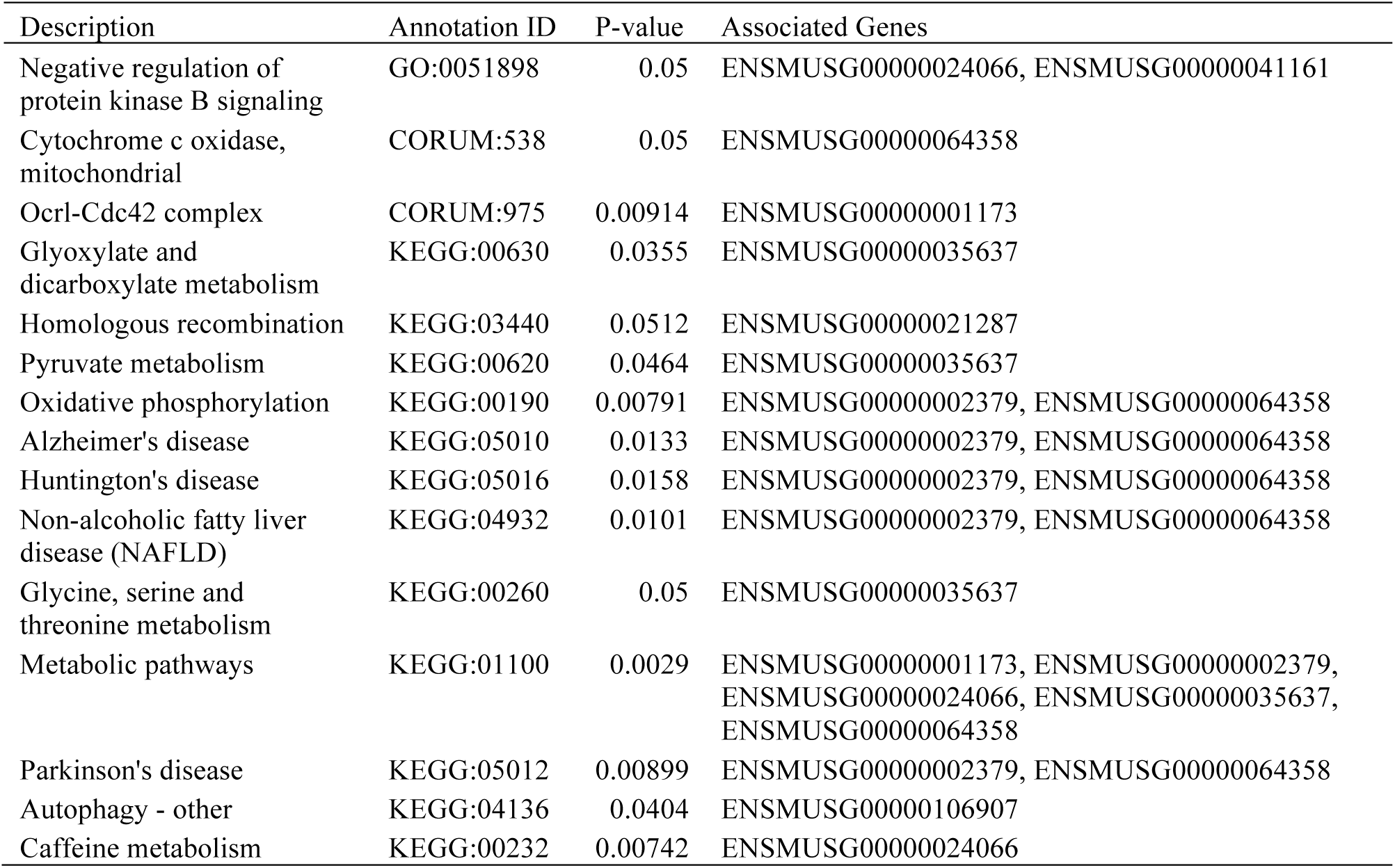
Overrepresented gene ontology (GO) terms from g:Profiler (q-value < 0.05) for the 19 outlier loci from tests for both selection and GEA. Associated genes shows which ensemble gene homologs from Table 2 are associated with each overrepresented term.

## DISCUSSION

In this study, we investigated patterns of divergent positive selection between urban and rural populations of *P. leucopus*, and identified significant associations between outlier SNPs and environmental variables relevant to urbanization. The majority of candidate loci were annotated with GO terms that are significantly associated with dietary metabolism, particularly breakdown of lipids and carbohydrates. We discuss what these findings mean for organisms inhabiting novel urban ecosystems, and more generally for understanding the ecological processes and time frame of local adaptation in changing environments.

Our previous study investigated non-synonymous polymorphisms in pooled transcriptome samples and we reported evidence for positive selection in genes dealing with metabolism, immunity, and methylation in NYC white-footed mice (Harris *et al.* 2013). This current study supports the phenotypic traits likely under selection in urban environments, identifying outlier genes that play major roles in metabolism, and to a lesser extent, immunity, but few outlier genes were identified in both the current and previous studies. The dataset analyzed here was much larger, included more sampling sites, and changed the inclusion criteria for outlier genes by using analyses that identify more recent signatures of selection, as opposed to longer-term evolutionary changes in non-synonymous substitutions. However, it is important to note that our study is still relatively small, including only six populations and eight individuals from each population. Increasing the number of individuals and sampling sites, especially including multiple cities as replicates, would likely greatly improve the associations found between environmental variables and allele frequencies (Lotterhos & Whitlock 2015). The latter approach may be unlikely, however, with each urban setting presenting a unique set of selective pressures leading to local adaptive responses, as shown with coat coloration in beach mice (*Peromyscus polionotus*) (Hoekstra *et al.* 2006) and climate related adaptation in the flowering plant (*Arabidopsis halleri)* (Rellstab *et al.* 2017). Despite potential issues with sample size, we did find two of the eleven previously identified candidate genes (Harris *et al.* 2013) to be direct matches to outliers in this current analysis (Serine protease inhibitor a3c and Solute carrier organic anion transporter 1A5), and two other genes were from the same gene families or involved in the same biological processes. One gene, an aldo-keto-reductase protein, is part of the same gene family as the aflatoxin reductase gene (Contig 10636-348) identified in this study. The aldo-keto reductase gene family comprises a large group of essential enzymes for metabolizing natural and foreign substances (Hyndman *et al.* 2003). The other is a cytochrome P450 (CYPA1A) gene involved in metabolism of drugs and lipids. *Peromyscus* directly express CYPA1A and Hsp90 (outlier from current SweeD analysis) when exposed to environmental toxins (Settachan 2001).

### Population genomics summary statistics

Before performing outlier tests, we initially calculated per-site nucleotide diversity and Tajima’s D. The Tajima’s D statistic was calculated per contig for each population. We found nucleotide diversity to be lower in all urban population compared to rural populations, supporting previous work that found a negative association between genome-wide SNP diversity and urbanization. That study included the six populations studied here and an additional 18 populations distributed along an urban-to-rural gradient (Munshi-South *et al.* 2016). While loss of genetic variation will reduce evolutionary potential and decrease the probability of local adaptation, selection may still act if adequate variation is present and genetic drift is not too strong (Donihue & Lambert 2015; Munshi-South *et al.* 2016). Tajima’s D is often used to identify signatures of selection, comparing observed to expected heterozygosity. For all our populations, Tajima’s D skewed positive, possibly explained by balancing selection. While balancing selection has been found to maintain variation in immune loci in fragmented urban population of bobcats (*Lynx rufus*) (Serieys *et al.* 2015), it is difficult to distinguish whether demography or selection drives Tajima’s D values in many cases (MacManes & Eisen 2014).

We have estimated the complex demographic history for *P. leucopus* populations in NYC (Harris *et al.* 2016), suggesting Tajima’s D may not be the best tool for identifying selection in this system. Outlier tests are more robust to demography and we explicitly accounted for the specific demographic history of *P. leucopus* in the null models used during analysis of our genome scan methods.

### Signatures of selection in urban populations from genome-wide scans

Over the past decade, genome scans have become feasible methods to detect and disentangle neutral and adaptive evolutionary processes for non-model organisms (De Villemereuil *et al.* 2014; Hoban *et al.* 2016). One method, BayeScan (Foll & Gaggiotti 2008), calculates the posterior probability that a site is under the influence of selection by testing models with and without selection. While BayeScan is relatively robust to confounding demographic processes (Pérez-Figueroa *et al.* 2010; De Villemereuil *et al.* 2014), population bottlenecks, hierarchical structure, recent migration, or variable times to most-recent-common-ancestor (MRCA) between populations can artificially inflate *F*_*ST*_ values (Hermisson 2009; Lotterhos & Whitlock 2014) and may still impact BayeScan (Savolainen *et al.* 2013; Lotterhos & Whitlock 2014). We minimized false positives by incorporating population structure and a specific demographic history for *P. leucopus* in NYC directly into the null distribution of *F*_*ST*_ (Harris *et al.* 2016). We only included outliers if their posterior probability was greater than probabilities calculated from these simulations. The outliers from BayeScan comprised 0.024% of the total number of loci analyzed from our RNASeq dataset, and 0.036% of the total loci using BayPass. These percentages are in line with candidates uncovered from a similar study (0.05%) that looked at high and low altitude populations of the plant *Senecio chrysanthemifolius* (Chapman *et al.* 2013). Many studies find higher percentages of outlier loci using BayeScan; for example, 4.5% in the American pika across its range in British Columbia (Henry & Russello 2013), and 5.7% in Atlantic herring across their range (Limborg *et al.* 2012). Our lower overall percentage of outliers may be due to differences in species or datasets between studies (false positive rate, power, sampling, genome size and composition are all variables that influence numbers of SNPs), or alternatively because of relatively recent isolation or moderate to weak selection in urban populations.

SweeD, another genome scan approach, examines patterns within a population’s SFS rather than allelic differentiation between populations. The main footprint that selective sweeps leave on the SFS is an excess of low- and high-frequency variants (Nielsen 2005). The SweepFinder method (Nielsen *et al.* 2005), recently upgraded to the NGS compatible SweeD (Pavlidis *et al.* 2013), uses a CLR test based on the ratio between the likelihood of a neutral and selective sweep hypothesis. As above, the weakness of hitchhiking methods is the confounding influence certain demographic processes have on the SFS (Hermisson 2009). However, building a robustly inferred demographic history into the null model substantially reduces false positive rates (Pavlidis *et al.* 2013). We included the *P. leucopus* demographic history into our analysis, and found 0.019% of the sequenced loci to contain SFS patterns indicative of selective sweeps. This rate is in line with other studies that reported that 0.5% of regions in domesticated rice (Wang *et al.* 2014), 0.02% of loci in black cottonwood (Zhou *et al.* 2014), and 0.02% of the gorilla genome (McManus *et al.* 2014) show evidence of selective sweeps or hitchhiking.

Several studies have shown that identifying outliers with multiple tests and diverse theoretical approaches is the best way to reduce false positives in genome outlier analyses (Nielsen 2005; Grossman *et al.* 2010; Hohenlohe *et al.* 2010b). We required candidate genes to show a signature of diversifying selection or a signature of a selective sweep, and they had to be associated with an environmental variable. We found several outliers identified in both BayeScan and BayPass (Table S2), however, there was no overlap between BayeScan / BayPass and SweeD outliers. This discrepancy is likely due to the different selection scenarios underlying each test, i.e. divergent local selection versus population-wide positive selection in the form of selective sweeps (Hermisson 2009). *F*_*ST*_ based methods respond to allelic divergence relatively quickly, while models for selective sweeps typically require nearly-fixed derived alleles (Hohenlohe *et al.* 2010b). Given the recent history of urbanization in NYC, many selective sweeps may be ongoing or otherwise incomplete. Selection may also be acting on standing genetic variation in the form of soft sweeps (Hermisson & Pennings 2005) that are not readily identified by SweeD.

### Environmental associations strengthen evidence of local adaptation to urbanization

GEA tests are a growing class of methods that identify loci that are associated with environmental factors (Joost *et al.* 2007; Coop *et al.* 2010; Frichot *et al.* 2013), and by accounting for underlying correlation structure of allele frequencies, may often be more powerful than traditional outlier tests (Savolainen *et al.* 2013). GEA tests come from the field of landscape genomics which incorporates tools from landscape genetics and population genomics to examine the effects of demography, migration, and selection, and ultimately identify local adaptation (Sork *et al.* 2013; Rellstab *et al.* 2015). Here we used LFMM (Frichot *et al.* 2013) and the AUX covariate model from BayPass on the full SNP dataset with environmental metrics of urbanization. LFMM performs better than other methods in the presence of hierarchical structure and when polygenic selection is acting on many loci with small effect (De Villemereuil *et al.* 2014). Hierarchical structure in our dataset includes urban and rural differentiation (Harris *et al.* 2015; Harris *et al.* 2016), patterns of geographic structure between mainland mice and Long Island, NY (Harris *et al.* 2016), and population structure between individual urban parks (Munshi-South & Kharchenko 2010). Simulations also suggest that LFMM is superior when sample size is less than 10 individuals per population, there is no pattern of IBD, and the study compares environmentally divergent habitats (Lotterhos & Whitlock 2015). We sampled eight white-footed mice per population, found no evidence of IBD (Munshi-South *et al.* 2016), and sampled environmentally divergent rural and urban locations.

Using GEA tests implemented in BayPass and LFMM, we found that 17 (12 %) and 4 (2.8 %) outliers, respectively, were significantly associated with one or more urbanization variables. These results are lower than other studies combining genome scans and GEA tests. Limborg *et al.* (2012) found 62.5% of the outliers identified in BayeScan were correlated with temperature or salinity in Atlantic herring, and 26.3% of genome scan outliers were associated with temperature or latitude in a tree species (De Kort *et al.* 2014). The lower overlap found in our study is likely due to the difficult nature in quantifying urbanization. Percent impervious surface, human population density, or binary classification as urban versus rural may not capture the specific, causative selection pressures acting on white-footed mouse populations (See Table S5 for environmental data). We used these metrics as general proxies for changing ecological processes in urbanized habitats. The percent of impervious surface around a park is likely representative of habitat fragmentation, as urban infrastructure changes the net primary productivity due to increasing percentages of impervious surface or artificial landscapes, parks and yards (Shochat *et al.* 2006). This fragmentation then leads to changing species interactions as migration is impeded or organisms are forced into smaller areas (Shochat *et al.* 2006). The percent human density surrounding an urban park can serve as a proxy for the multitude of ecological changes humans impose on their surrounding environment. Urbanization and increasing human density change the types and availability of resources in the altered habitat (McKinney 2002; Sih *et al.* 2011). Finally, classifying our sites as urban or rural can generally capture the main differences in urban and natural sites. For example, pollution is a major consequence of urbanization (Donihue & Lambert 2015), and urban areas often include increased chemical, noise, or light pollution (Sih *et al.* 2011).

Between divergent allele frequencies, a skewed SFS, environmental associations, and overrepresented GO terms, we find several overlapping lines of evidence that support rapid divergent selection in white-footed mice. Our results support the growing body of evidence (Donihue & Lambert 2015) that finds urbanization directly impacts the ecology and evolution of species. However, to fully support the hypothesis that organisms adapt to urban habitats, it is still necessary to link genetic changes to measurable phenotypic differences and measure direct fitness benefits. Past urban evolutionary studies often focus solely on phenotypic (Yeh 2004; Partecke *et al.* 2006; Cheptou *et al.* 2008; Thompson *et al.* 2016) or genetic (Wandeler *et al.* 2003; Noë l & Lapointe 2010; Mueller *et al.* 2013; Lourenco *et al.* 2017) differences between populations in and outside of cities. However, researchers are beginning to examine both the genotype and phenotype in parallel instances of urban evolution (Whitehead *et al.* 2010; Wirgin *et al.* 2011; Hof *et al.* 2016), which is key to understanding how urbanization affects the evolution of species. In the future, the gene annotations for our predicted outlier genes can help determine which phenotypic traits to measure in urban *P. leucopus* populations.

### Functional roles of candidate genes: quality of urban diet?

The model rodents *Mus musculus, Rattus norvegicus,* and *Cricetulus griseus* all have deeply sequenced, assembled and annotated reference genomes. These resources allowed us to annotate 89.5% of outlier loci with high quality functional information. Urban *P. leucopus* exhibited signatures of positive selection in genes with GO terms overrepresented for organismal metabolic processes, specifically digestion and metabolism of lipids and carbohydrates.

Mitochondrial genes identified as outliers (Table S2) were largely responsible for the overrepresentation of metabolic process. While we can only speculate until further physiological studies are conducted, our evidence suggests that the evolution of mitochondrial and metabolic processes has been important to the success of *P. leucopus* living in NYC’s urban forests. Mitochondrial genes have often been used to describe neutral population variation, but researchers have found ample evidence of selection acting on the mitochondrial genome (Oliveira *et al.* 2008; Balloux 2010). For example, specific mitochondrial haplotypes are associated with more efficient thermogenesis and higher fitness in over-wintering shrews (Fontanillas *et al.* 2005). Pergams & Lacy (2007) found complete mitochondrial haplotype replacement in contemporary *P. leucopus* in Chicago compared to haplotypes sequenced from museum skins collected before urbanization. The agent of selection is not clear, but Munshi-South and Nagy (2014) also identified signatures of selection (or alternatively population expansion) in mitochondrial D-loop haplotypes from contemporary *P. leucopus* in NYC. Many mitochondrial functions are affected by the same environmental variables that change in response to urbanization, such as temperature (Balloux 2010), reduced migration (Lankau & Strauss 2011; Munshi-South 2012), or resource availability (Burcelin *et al.* 2002).

Urban *P. leucopus* may experience different energy budgets, physiological stressors or diets compared to rural counterparts. We found a substantial number of candidate genes with functions related to the metabolism and transport of lipids and carbohydrates, and the most common overrepresented GO terms involved lipid metabolism and homeostasis (Table S4). In the full outlier analysis, two genes are particularly interesting as targets of diet-mediated selection. The first gene, *FADS1,* is a fatty acid desaturase important for the biosynthesis of omega-3 and -6 fatty acids (long-chain polyunsaturated fatty acids, LCPUFA) from plant sources. Recent evidence suggests that the *FADS* gene family has been an important target of selection in humans during the transition from hunter-gather to agricultural societies (Ye *et al.* 2017). Alleles linked to upregulated biosynthesis of LCPUFAs (naturally low in plant based diets) increased in frequency after the Neolithic Revolution (Ye *et al.* 2017). We aligned our homologous *FADS1* contig with human transcripts to identify whether *P. leucopus* had any relevant alleles, but our sequenced populations did not contain SNPs at any relevant loci. The list of outliers also contained *APOB-100*, which is the primary apolipoprotein that binds and transports lipids, including both forms of cholesterol (HDL and LDL).

When we investigated only candidate genes that were identified by both an outlier test and GEA test, we found similar patterns suggesting *P. leucopus* in urban environments may be adapting to novel food resources. These genes were strongly correlated with environmental measures of urbanization, with clearly divergent allele frequencies between urban and rural sites (Fig. 3B), suggesting that selection is acting on standing genetic variation in urban environments. The most significant overrepresented GO term involved regulation of protein kinase B (*AKT*). *AKT* is a key molecule in the insulin signaling pathway, important for promoting glucose storage and regulating glucose in the bloodstream between fed and fasting states (Boucher *et al.* 2014). Glycine metabolism was also overrepresented; increased amounts of glycine may be important for regulating high-fat, high-sugar diets by decreasing concentrations of free fatty acids and triglycerides (Wang *et al.* 2013). Finally, our candidate list contained genes significantly associated with non-alcoholic fatty liver disease (NAFLD). NAFLD is a major hallmark of obesity and diabetes and can be induced through increased uptake of dietary fatty acids (Fabbrini *et al.* 2010).

These candidate genes suggest that white-footed mice in isolated urban parks may be evolving in response to food resource differences between urban and rural habitats. This finding is corroborated by recent evidence that urban white-footed mice in NYC have shorter upper and lower tooth rows than rural mice (Yu *et al.* 2017). Lower quality food in the diet often requires increased chewing and is accompanied with larger occlusal surfaces, and subsequently, longer toothrows (Ungar 2010). One prediction is that urban *P. leucopus* consume a diet with a substantially higher fat content than diets of rural populations. The typical diet of *P. leucopus* across its range consists of arthropods, fruits, nuts, various green vegetation, and fungus (Wolff *et al.* 1985). Given that white-footed mice are opportunistic generalists, many different food resources could differ between urban and rural habitats. Urbanization in NYC has produced relatively small green patches that are surrounded by a dense urban matrix, and *P. leucopus* in NYC may successfully take advantage of invasive plant species, different arthropod communities, or increased human food waste in and around their urban habitats. Local adaptation in urban populations may allow these mice to more efficiently metabolize different types or amounts of lipids and carbohydrates, although field studies are needed to examine the link between these genetic changes and diet in NYC.

## ACKNOWLEDGMENTS

We thank Mike Hickerson for his helpful comments and advice on many analyses and for access to lab space for analyses and writing. We thank Diego Alvarado-Serrano, Alexander T. Xue, Tyler Joseph, and Champak Reddy for their invaluable comments and advice concerning bioinformatics and demographic analyses. Three anonymous reviewers and Prof. Stuart B. Piertney also provided very helpful comments on the manuscript through Axios Review, as did Dr. Aurélie Bonin and four anonymous reviewers for this journal. This research was supported by the National Institute of General Medical Sciences of the National Institutes of Health under award number R15GM099055 to JM-S and a NSF Graduate Research Fellowship to SEH. The content is solely the responsibility of the authors and does not represent the official views of the National Institutes of Health.

## Data Accessibility

-VCF file of SNP genotypes used for demographic inference: Dryad doi:10.5061/dryad.d48f9

-raw sequencing files for transcriptome data: GenBank Sequence Read Archive (SRA Accession no. <u>SRP020005</u>)

-Transcriptome contigs: Dryad doi:<u>10.5061/dryad.6hc0f</u>

## SUPPORTING INFORMATION

**Table S1.** Average pairwise *F*_*ST*_ among six *P. leucopus* populations based on transcriptome-derived SNPs.

**Table S2.** Excel file containing the full list of outlier contigs (N = 381), the outlier SNP position, and test(s) that identified the outlier SNP. Remaining columns list the homologous Ensemble *Mus musculus* gene ID and name.

**Table S3.** Excel file containing results from g:Profiler for overrepresented GO terms from the full list of outlier contigs in Table S2. The table also includes the homologous *Mus musculus* genes that are associated with each GO term.

**Table S4.** Excel file containing Revigo results. Enriched GO terms from g:Profiler are sorted into largest parent terms and listed based on the frequency of occurrence.

**Table S5.** Environmental variable values for each individual mouse. Impervious = mean % impervious surface in 2 km buffer. Density = human population per 2 km buffer. Urban or Rural = classification as urban or rural site.

## REFERENCES

Aken BL, Ayling S, Barrell D et al.(2016) The Ensembl gene annotation system. Database, 2016, baw093.

Balloux F,(2010) The worm in the fruit of the mitochondrial DNA tree. Heredity, 104, 419–420.

Barrett RDH, Schluter D (2008) Adaptation from standing genetic variation. Trends in Ecology & Evolution, 23, 38–44.

Bonin A (2008) Population genomics: a new generation of genome scans to bridge the gap with functional genomics. Molecular Ecology, 17, 3583–4.

Boucher J, Kleinridders A, Kahn CR (2014) Insulin Receptor Signaling in Normal and Insulin-Resistant States. Cold Spring Harbor Perspectives in Biology, 6, a009191–a009191.

Burcelin R, Crivelli V, Dacosta A, Roy-Tirelli A, Thorens B (2002) Heterogeneous metabolic adaptation of C57BL/6J mice to high-fat diet. American Journal of Physiology Endocrinology and Metabolism, 282, E834–E842.

Chace JF, Walsh JJ (2004) Urban effects on native avifauna: a review. Landscape and Urban Planning, 74, 46–69.

Chapman MA, Hiscock SJ, Filatov DA (2013) Genomic Divergence during Speciation Driven by Adaptation to Altitude. Molecular Biology and Evolution, 30, 2553–67.

Cheptou P-O, Carrue O, Rouifed S, Cantarel A (2008) Rapid evolution of seed dispersal in an urban environment in the weed Crepis sancta. Proceedings of the National Academy of Sciences of the United States of America, 105, 3796–9.

Conesa A, Gö tz S, García-Gómez JM et al.(2005) Blast2GO: a universal tool for annotation, visualization and analysis in functional genomics research. Bioinformatics, 21, 3674–6.

Coop G, Witonsky D, Di Rienzo A, Pritchard JK (2010) Using environmental correlations to identify loci underlying local adaptation. Genetics, 185, 1411–23.

Danecek P, Auton A, Abecasis G et al.(2011) The variant call format and VCFtools. Bioinformatics, 27, 2156–2158.

DePristo MA, Banks E, Poplin R et al.(2011) A framework for variation discovery and genotyping using next-generation DNA sequencing data. Nature Genetics, 43, 491–8.

Donihue CM, Lambert MR (2015) Adaptive evolution in urban ecosystems. AMBIO, 44, 194–203.

Ellison CE, Hall C, Kowbel D et al.(2011) Population genomics and local adaptation in wild isolates of a model microbial eukaryote. Proceedings of the National Academy of Sciences, 108, 2831–2836.

Excoffier L, Dupanloup I, Huerta-Sanchez E, Sousa VC, Foll M (2013) Robust Demographic Inference from Genomic and SNP Data. PLoS Genetics, 9, e1003905.

Excoffier L, Hofer T, Foll M (2009) Detecting loci under selection in a hierarchically structured population. Heredity, 103, 285–98.

Fabbrini E, Sullivan S, Klein S (2010) Obesity and nonalcoholic fatty liver disease: Biochemical, metabolic, and clinical implications. Hepatology, 51, 679–689.

Foll M, Gaggiotti O (2008) A genome-scan method to identify selected loci appropriate for both dominant and codominant markers: a Bayesian perspective. Genetics, 180, 977–93.

Fontanillas P, Dépraz A, Giorgi MS, Perrin N (2005) Nonshivering thermogenesis capacity associated to mitochondrial DNA haplotypes and gender in the greater white-toothed shrew, Crocidura russula. Molecular Ecology, 14, 661–670.

Frichot E, François O (2015) LEA: An R package for landscape and ecological association studies. Methods in Ecology and Evolution, 6.

Frichot E, Schoville SD, Bouchard G, François O (2013) Testing for associations between loci and environmental gradients using latent factor mixed models. Molecular Biology andEvolution, 30, 1687–1699.

Gautier M (2015) Genome-wide scan for adaptive divergence and association with population-specific covariates. Genetics, 201, 1555–1579.

Götz S, García-Gómez JM, Terol J et al.(2008) High-throughput functional annotation and data mining with the Blast2GO suite. Nucleic Acids Research, 36, 3420–35.

Grossman SR, Shylakhter I, Karlsson EK et al.(2010) A composite of multiple signals distinguishes causal variants in regions of positive selection. Science, 327, 883–6.

Gutenkunst RN, Hernandez RD, Williamson SH, Bustamante CD (2009) Inferring the joint demographic history of multiple populations from multidimensional SNP frequency data. PLoS Genetics, 5, e1000695.

Harris SE, Munshi-South J, Obergfell C, O,Neill R (2013) Signatures of Rapid Evolution in Urban and Rural Transcriptomes of White-Footed Mice (Peromyscus leucopus) in the New York Metropolitan Area. PLoS ONE, 8, e74938.

Harris, SE O ,Neill RJ, Munshi-South J (2015) Transcriptome resources for the white-footed mouse (Peromyscus leucopus): new genomic tools for investigating ecologically divergent urban and rural populations. Molecular Ecology Resources, 15, 382–394.

Harris SE, Xue AT, Alvarado-Serrano D et al.(2016) Urbanization shapes the demographic history of a native rodent (the white-footed mouse, Peromyscus leucopus) in New York City. Biology Letters, 12, 20150983-.

Henry P, Russello MA (2013) Adaptive divergence along environmental gradients in a climate-change-sensitive mammal. Ecology and Evolution, 3, 3906–3917.

Hermisson J (2009) Who believes in whole-genome scans for selection? Heredity, 103, 283–284.

Hermisson J, Pennings PS (2005) Soft sweeps: Molecular population genetics of adaptation fromstanding genetic variation. Genetics, 169, 2335–2352.

Hoban S, Kelley JL, Lotterhos KE et al.(2016) Finding the Genomic Basis of Local Adaptation: Pitfalls, Practical Solutions, and Future Directions. The American Naturalist, 188, 379–397.

Hoekstra HE, Hirschmann RJ, Bundey RA, Insel PA, Crossland JP (2006) A Single Amino Acid Mutation Contributes to Adaptive Beach Mouse Color Pattern. Science, 313, 101–104.

Hof AEV, Campagne P, Rigden DJ et al.(2016) The industrial melanism mutation in British peppered moths is a transposable element. Nature, 534, 102–105.

Hohenlohe PA, Bassham S, Etter PD et al.(2010a) Population genomics of parallel adaptation in threespine stickleback using sequenced RAD tags. PLoS Genetics, 6, e1000862.

Hohenlohe PA, Phillips PC, Cresko WA (2010b) Using Population Genomics to Detect Selection in Natural Populations: Key Concepts and Methodological Considerations. International Journal of Plant Sciences, 171, 1059–1071.

Hyndman D, Bauman DR, Heredia V V., Penning TM (2003) The aldo-keto reductase superfamily homepage. Chemico-Biological Interactions, 143–144, 621–631.

Joost S, Bonin A, Bruford MW et al.(2007) A spatial analysis method (SAM) to detect candidate loci for selection: towards a landscape genomics approach to adaptation. Molecular Ecology, 16, 3955–69.

Kanehisa M, Goto S, Sato Y et al.(2014) Data, information, knowledge and principle: Back to metabolism in KEGG. Nucleic Acids Research, 42, 199–205.

Kim Y, Stephan W (2002) Detecting a local signature of genetic hitchhiking along a recombining chromosome. Genetics, 160, 765–777.

De Kort H, Vandepitte K, Bruun HH et al.(2014) Landscape genomics and a common garden trial reveal adaptive differentiation to temperature across Europe in the tree species Alnusglutinosa. Molecular Ecology, 4709–4721.

Lankau RA, Strauss SY (2011) Newly rare or newly common: evolutionary feedbacks through changes in population density and relative species abundance, and their management implications. Evolutionary Applications, 4, 338–353.

Li J, Li H, Jakobsson M et al.(2012) Joint analysis of demography and selection in population genetics: where do we stand and where could we go? Molecular Ecology, 28, 28–44.

Limborg MT, Helyar SJ, De Bruyn M et al.(2012) Environmental selection on transcriptome-derived SNPs in a high gene flow marine fish, the Atlantic herring (Clupea harengus). Molecular ecology, 21, 3686–703.

Linnen CR, Kingsley EP, Jensen JD, Hoekstra HE (2009) On the origin and spread of an adaptive allele in deer mice. Science, 325, 1095–8.

Lotterhos KE, Whitlock MC (2014) Evaluation of demographic history and neutral parameterization on the performance of F ST outlier tests. Molecular Ecology, 23, 2178– 2192.

Lotterhos KE, Whitlock MC (2015) The relative power of genome scans to detect local adaptation depends on sampling design and statistical method. Molecular Ecology, 24, 1031–1046.

Lourenco A, Alvarez D, Wang IJ, Velo-Anton G (2017) Trapped within the city: integrating demography, time since isolation and population-specific traits to assess the genetic effects of urbanization. Molecular Ecology, 26, 1498–1514.

MacManes MD, Eisen MB (2014) Characterization of the transcriptome, nucleotide sequence polymorphism, and natural selection in the desert adapted mouse Peromyscus eremicus. PeerJ, 2, e642.

McKinney ML (2002) Urbanization, biodiversity, and conservation. Bioscience, 52, 883–890.

McManus KF, Kelley JL, Song S et al.(2014) Inference of Gorilla Demographic and Selective History from Whole-Genome Sequence Data. Molecular Biology and Evolution, 32, 600–612.

Merilä J, Hendry AP (2014) Climate change, adaptation, and phenotypic plasticity: the problem and the evidence. Evolutionary Applications, 7, 1–14.

Mueller JC, Partecke J, Hatchwell BJ, Gaston KJ, Evans KL (2013) Candidate gene polymorphisms for behavioural adaptations during urbanization in blackbirds. Molecular Ecology, 22, 3629–3637.

Mullen LM, Hoekstra HE (2008) Natural selection along an environmental gradient: a classic cline in mouse pigmentation. Evolution, 62, 1555–70.

Munshi-South J (2012) Urban landscape genetics: canopy cover predicts gene flow between white-footed mouse (Peromyscus leucopus) populations in New York City. Molecular Ecology, 21, 1360–1378.

Munshi-South J, Kharchenko K (2010) Rapid, pervasive genetic differentiation of urban white-footed mouse (Peromyscus leucopus) populations in New York City. Molecular Ecology, 19, 4242–4254.

Munshi-South J, Nagy C (2014) Urban park characteristics, genetic variation, and historical demography of white-footed mouse ( Peromyscus leucopus) populations in New York City. PeerJ, 2, e310.

Munshi-South J, Richardson JL (2017) Peromyscus transcriptomics: Understanding adaptation and gene expression plasticity within and between species of deer mice. Seminars in Cell & Developmental Biology, 61, 131–139.

Munshi-South J, Zolnik CP, Harris SE (2016) Population genomics of the Anthropocene: urbani zation is negatively associated with genome-wide variation in white-footed mouse populations. Evolutionary Applications, doi:10.1111/eva.12357.

Natarajan C, Inoguchi N, Weber RE et al.(2013) Epistasis Among Adaptive Mutations in Deer Mouse Hemoglobin. Science, 340, 1324–1327.

Nielsen R (2005) Molecular signatures of natural selection. Annual Review of Genetics, 39, 197–218.

Nielsen R, Williamson S, Kim Y et al.(2005) Genomic scans for selective sweeps using SNP data. Genome research, 15, 1566–75.

Noë l S, Lapointe F (2010) Urban conservation genetics: Study of a terrestrial salamander in the city., 143, 2823–2831.

Ogata H, Goto S, Sato K et al.(1999) KEGG: Kyoto encyclopedia of genes and genomes. Nucleic Acids Research, 27, 29–34.

Oleksyk TK, Smith MW, O ,Brien SJ (2010) Genome-wide scans for footprints of natural selection. Philosophical transactions of the Royal Society of London. Series B, Biological sciences, 365, 185–205.

Oliveira DCSG, Raychoudhury R, Lavrov D V, Werren JH (2008) Rapidly evolving mitochondrial genome and directional selection in mitochondrial genes in the parasitic wasp Nasonia (Hymenoptera: Pteromalidae). Molecular Biology and Evolution, 25, 2167–2180.

Orr HA (2005) The genetic theory of adaptation: a brief history. Nature Reviews Genetics, 6, 119–27.

Partecke J, Schwabl I, Gwinner E (2006) Stress and the city: Urbanization and its effects on the stress physiology in European Blackbirds. Ecology, 87, 1945–1952.

Patterson N, Price AL, Reich D (2006) Population structure and eigenanalysis. PLoS Genetics, 2, e190.

Pavlidis P, Jensen JD, Stephan W (2010) Searching for Footprints of Positive Selection in Whole-genome SNP Data from Non-equilibrium Populations. Genetics.

Pavlidis P, Živkovic D, Stamatakis A, Alachiotis N (2013) SweeD: likelihood-based detection of selective sweeps in thousands of genomes. Molecular Biology and Evolution, 30, 2224–34.

Pérez-Figueroa A, García-Pereira MJ, Saura M, Rolán-Alvarez E, Caballero A (2010) Comparing three different methods to detect selective loci using dominant markers. Journal of Evolutionary Biology, 23, 2267–2276.

Pergams ORW, Lacy RC (2008) Rapid morphological and genetic change in Chicago-area Peromyscus. Molecular Ecology, 17, 450–63.

Reid NM, Proestou DA, Clark BW et al.(2016) The genomic landscape of rapid repeated evolutionary adaptation to toxic pollution in wild fish. Science, 354, 1305–1308.

Reimand J, Arak T, Adler P et al.(2016) g:Profiler—a web server for functional interpretation of gene lists (2016 update). Nucleic Acids Research, 44, W83–W89.

Rellstab C, Fischer MC, Zoller S et al.(2017) Local adaptation (mostly) remains local: reassessing environmental associations of climate-related candidate SNPs in Arabidopsis halleri. Heredity, 118, 193–201.

Rellstab C, Gugerli F, Eckert AJ, Hancock AM, Holderegger R (2015) A practical guide to environmental association analysis in landscape genomics. Molecular Ecology, 24, 4348–4370.

Rockman M (2012) The QTN program and the alleles that matter for evolution: all that,s gold does not glitter. Evolution, 66, 1–17.

Rogic A, Tessier N, Legendre P, Lapointe F-J, Millien V (2013) Genetic structure of the white-footed mouse in the context of the emergence of Lyme disease in southern Québec. Ecology and Evolution, 3, 2075–88.

Rytwinski T, Fahrig L (2007) Effect of road density on abundance of white-footed mice. Landscape Ecology, 22, 1501–1512.

Savolainen O, Lascoux M, Merilä J (2013) Ecological genomics of local adaptation. Nature Reviews Genetics, 14, 807–20.

Serieys LEK, Lea A, Pollinger JP, Riley SPD, Wayne RK (2015) Disease and freeways drive genetic change in urban bobcat populations. Evolutionary Applications, 8, 75–92.

Settachan D (2001) Mechanistic and molecular studies into the effects of 2,3,7,8-tetrachlorodibenzo-p-dioxin and similar compounds in the deer mouse, Peromyscus maniculatus. Texas Tech University.

Shochat E, Warren PS, Faeth SH, McIntyre NE, Hope D (2006) From patterns to emerging processes in mechanistic urban ecology. Trends in Ecology and Evolution, 21, 186–91.

Sih A, Ferrari MCO, Harris DJ (2011) Evolution and behavioural responses to human-induced rapid environmental change. Evolutionary Applications, 4, 367–387.

Sork VL, Aitken SN, Dyer RJ et al.(2013) Putting the landscape into the genomics of trees: Approaches for understanding local adaptation and population responses to changing climate. Tree Genetics and Genomes, 9, 901–911.

Stapley J, Reger J, Feulner PGD et al.(2010) Adaptation Genomics: the next generation. Trends in Ecology & Evolution, 25, 705–712.

Stinchcombe JR, Hoekstra HE (2008) Combining population genomics and quantitative genetics: finding the genes underlying ecologically important traits. Heredity, 100, 158–70.

Storz JF, Runck AM, Moriyama H, Weber RE, Fago A (2010) Genetic differences in hemoglobin function between highland and lowland deer mice. The Journal of Experimental Biology, 213, 2565–74.

Storz JF, Runck AM, Sabatino SJ et al.(2009) Evolutionary and functional insights into the mechanism underlying high-altitude adaptation of deer mouse hemoglobin. Proceedings of the National Academy of Sciences of the United States of America, 106, 14450–5.

Storz J, Sabatino S, Hoffmann F (2007) The molecular basis of high-altitude adaptation in deer mice. PLoS Genetics, 3.

Supek F, Bosnjak M, Skunca N, Smuc T (2011) Revigo summarizes and visualizes long lists of gene ontology terms. PLoS ONE, 6.

Tajima F (1983) Evolutionary relationship of DNA sequences in finite populations. Genetics, 105, 437–460.

Tajima F (1989) Statistical method for testing the neutral mutation hypothesis by DNA polymorphism. Genetics, 123, 585–95.

Thompson KA, Renaudin M, Johnson MTJ (2016) Urbanization drives the evolution of parallel clines in plant populations. Proceedings of the Royal Society B: Biological Sciences, 283, 20162180.

Tiffin P, Ross-Ibarra J (2014) Advances and limits of using population genetics to understand local adaptation. Trends in Ecology & Evolution, 29, 673–680.

Tomassini A, Colangelo P, Agnelli P et al.(2014) Cranial size has increased over 133 years in a common bat, Pipistrellus kuhlii : a response to changing climate or urbanization? Journal Biogeography, 41, 944–953.

Turner TL, Bourne EC, Von Wettberg EJ, Hu TT, Nuzhdin S V (2010) Population resequencing reveals local adaptation of Arabidopsis lyrata to serpentine soils. Nature Genetics, 42, 260–3.

Ungar PS (2010) Mammal Teeth: Origin, Evolution, and Diversity. Johns Hopkins University Press, Baltimore, MD.

De Villemereuil P, Frichot É, Bazin É, François O, Gaggiotti OE (2014) Genome scan methods against more complex models: When and how much should we trust them? Molecular Ecology, 23, 2006–2019.

Vitti JJ, Grossman SR, Sabeti PC (2013) Detecting natural selection in genomic data. Annual review of genetics, 47, 97–120.

Wandeler P, Funk SM, Largiadèr CR, Gloor S, Breitenmoser U (2003) The city-fox phenomenon: genetic consequences of a recent colonization of urban habitat. Molecular Ecology, 12, 647–56.

Wang W, Wu Z, Dai Z et al.(2013) Glycine metabolism in animals and humans: Implications for nutrition and health. Amino Acids, 45, 463–477.

Wang M, Yu Y, Haberer G et al.(2014) The genome sequence of African rice (Oryza glaberrima) and evidence for independent domestication. Nature Genetics, 982–988.

Weber JN, Peterson BK, Hoekstra HE (2013) Discrete genetic modules are responsible for complex burrow evolution in Peromyscus mice. Nature, 493, 402–405.

Whitehead A, Triant D, Champlin D, Nacci D (2010) Comparative transcriptomics implicates mechanisms of evolved pollution tolerance in a killifish population. Molecular Ecology, 19, 5186–5203.

Wirgin I, Roy NK, Loftus M et al.(2011) Mechanistic basis of resistance to PCBs in Atlantic tomcod from the Hudson River. Science (New York, N.Y.), 331, 1322–5.

De Wit P, Palumbi SR (2013) Transcriptome-wide polymorphisms of red abalone (Haliotis rufescens) reveal patterns of gene flow and local adaptation. Molecular Ecology, 22, 2884–97.

Wolff JO, Dueser RD, Berry K (1985) Food Habits of Sympatric Peromyscus leucopus and Peromyscus maniculatus. Journal of Mammalogy, 66, 795–798.

Wright S (1951) The genetical structure of populations. Annals of Eugenics, 323–354.

Ye K, Gao F, Wang D, Bar-Yosef O, Keinan A (2017) Dietary adaptation of FADS genes in Europe varied across time and geography. Nature Ecology & Evolution, 1, 167.

Yeh PJ (2004) Rapid evolution of a sexually selected trait following population establishment in a novel habitat. Evolution, 58, 166–174.

Yu A, Munshi-south J, Sargis EJ et al.(2017) Morphological Differentiation in White-Footed Mouse ( Mammalia: Rodentia: Cricetidae: Peromyscus leucopus) Populations from the New York City Metropolitan Area. Bulletin of the Peabody Museum of Natural History, 58, 3–16.

Zhou L, Bawa R, Holliday JA (2014) Exome resequencing reveals signatures of demographic and adaptive processes across the genome and range of black cottonwood (Populus trichocarpa). Molecular Ecology, 23, 2486–2499.

